# Manganese uptake, mediated by SloABC and MntH, is essential for the fitness of *Streptococcus mutans*

**DOI:** 10.1101/817585

**Authors:** Jessica K. Kajfasz, Callahan Katrak, Tridib Ganguly, Jonathan Vargas, Logan Wright, Zachary T. Peters, Grace A. Spatafora, Jacqueline Abranches, José A. Lemos

## Abstract

Early epidemiological studies implicated manganese (Mn) as a possible caries-promoting agent while laboratory studies have indicated that manganese stimulates the expression of virulence-related factors in the dental pathogen *Streptococcus mutans*. To better understand the importance of manganese homeostasis to *S. mutans* pathophysiology, we first used RNA sequencing to obtain the global transcriptional profile of *S. mutans* UA159 grown under Mn-restricted conditions. Among the most highly expressed genes were the entire *sloABC* operon, encoding a dual iron/manganese transporter, and an uncharacterized gene, herein *mntH*, that codes for a protein bearing strong similarity to Nramp-type transporters. While inactivation of *sloC*, which encodes the lipoprotein receptor of the SloABC system, or *mntH* alone had no major consequence on the overall fitness of *S. mutans*, simultaneous inactivation of *sloC* and *mntH* (Δ*sloC*Δ*mntH*) impaired growth and survival under Mn-restricted conditions, including in human saliva or in the presence of calprotectin. Further, disruption of Mn transport resulted in diminished stress tolerance and reduced biofilm formation in the presence of sucrose. These phenotypes were markedly improved when cells were provided with excess Mn. Metal quantifications revealed that the single mutant strains contain similar intracellular levels of Mn as the parent strain, whereas Mn was nearly undetectable in the Δ*sloCΔmntH* strain. Collectively, these results reveal that SloABC and MntH work independently and cooperatively to promote cell growth under Mn-restricted conditions, and that mauitanence of Mn homeostasis is essential for the expression of major virulence attributes in *S. mutans*.

**IMPORTANCE:** As trace biometals such as manganese (Mn) are important for all forms of life, the ability to regulate biometals availability during infection is an essential trait of successful bacterial pathogens. Here, we showed that the caries pathogen *Streptococcus mutans* utilizes two Mn transport systems, namely SloABC and MntH, to acquire Mn from the environment, and that the ability to maintain the cellular levels of Mn is important for the manifestation of characteristics that associate *S. mutans* with dental caries. Our results indicate that the development of strategies to deprive *S. mutans* of Mn hold promise in the combat against this important bacterial pathogen.

## INTRODUCTION

Trace metals are essential for all domains of life by serving as structural and catalytic cofactors with approximately 50% of all enzymes in cells requiring a metal cofactor for proper function [1]. During microbial infections, the ability of the invading pathogen to acquire iron (Fe), manganese (Mn) and zinc (Zn) becomes particularly relevant as the host employs several mechanisms to sequester these essential biometals as part of an active response known as nutritional immunity [2–4]. Specifically, Fe-binding proteins such as transferrin (in serum) and lactoferrin (in secretions) are produced by the host to chelate Fe, thereby restricting its bioavailability to invading pathogens. Similarly, Mn and Zn are actively sequestered by calprotectin, a S100 family protein that constitutes about 60% of the total proteins in neutrophils [3]. To overcome this micronutrient limitation, bacteria evolved a number of mechanisms for metal acquisition, which include the production of low molecular weight molecules (metallophores) for extracellular metal capture, high-affinity membrane-associated metal transporters, as well as tools for the direct acquisition from host molecules and proteins (metal piracy) [4].

*Streptococcus mutans* is regarded as a keystone pathogen in dental caries due to its ability to change the oral biofilm architecture and environment such that it fosters the outgrowth of acidogenic and aciduric species (including *S. mutans* itself) at the expense of the commensal bacteria associated with oral health [5]. The cariogenic potential of *S. mutans* resides in its ability to (i) form robust biofilms on tooth surfaces in a sucrose-dependent manner, (ii) produce and tolerate large amounts of lactic acid, the major end product of its fermentative metabolism, and (iii) cope with oxidative stress that arises from the environmental reduction of oxygen and the production of hydrogen peroxide (H_2_O_2_) by competing neighbor species [6]. In addition to dental caries, *S. mutans* is also one of the causative agents of infective endocarditis, a life-threatening bacterial infection of the endocardium [7].

Previous studies conducted during the 1970s and 1980s have indicated a possible relationship between biometal availability in the oral cavity and caries incidence [8–12]. In particular, high rates of caries were linked to elevated levels of Mn in drinking water [8, 10, 12]. Despite the existence of conflicting clinical data questioning this correlation [9, 11], few studies have directly investigated the significance of Mn in the pathophysiology of oral streptococci [13–21]. An early study aiming to determine the trace element requirement of oral streptococci concluded that Mn was the only trace metal absolutely required for the growth of cariogenic and noncariogenic streptococci in the laboratory setting [18], a finding that was later confirmed by a second group of investigators [13]. In addition, Mn was shown to stimulate dextran-dependent aggregation in *Streptococcus criceti* (formerly *S. cricetus*) [22], a trait that was found to be mediated by surface-associated glucan-binding proteins (GBPs) and critical to sucrose-dependent adhesion and biofilm formation [23]. Subsequent studies using both *S. criceti* and *Streptococcus sobrinus* strains showed that metal chelating agents such as citrate or EDTA reversibly inhibit glucan-induced aggregation, thereby preventing sucrose-dependent adhesion [20]. In addition, confocal microscopy analysis of *S. mutans* UA159 biofilms grown in the presence of sucrose revealed that Mn-depleted biofilms formed large cell clumps that were more easily washed away than biofilms formed under Mn-replete conditions [14]. Finally, Mn was shown to stimulate carbohydrate metabolism in *S. mutans*, in particular the synthesis of glycogen-like intracellular polysaccharide (IPS) stores [17]. Finally, when added to drinking water, Mn was shown to increase the cariogenic potential of *S. mutans* in a germ-free rat model [17]. It should also be noted that Mn is known to play an important role in the oxidative stress responses of lactic acid bacteria by directly interacting with and scavenging superoxide radicals, by serving as the enzymatic co-factor of the superoxide dismutase enzyme, and by replacing Fe as an enzymatic co-factor, thereby protecting Fe-binding proteins from the irreversible damage of Fenton chemistry [24]. Collectively, the picture that emerges from these studies is that Mn may serve as a caries-promoting agent by stimulating bacterial metabolism, facilitating sucrose-dependent biofilm formation and, possibly, by conferring protection against the oxidative stresses encountered in dental plaque.

Because the nutrients available in the oral cavity derive, in large part, from the diet, the concentration of Mn in human saliva has been shown to fluctuate from as little as 1 μM [9, 11] to as high as 36 μM [25]. Taking into consideration that Mn is restricted to the nanomolar range in plasma [26], the concentration of Mn in saliva is unlikely to be a growth-limiting factor for most oral bacteria. Yet, fluctuations in Mn levels may serve as a cue for *S. mutans* to sense the environment and adjust its metabolism accordingly by favoring a biofilm survival mode over an active growth and/or dispersion mode. Beyond the oral environment, the ability to scavenge Mn in environments in which this metal is known to be restricted, such as the bloodstream and internal organs, has proven to be an essential trait for bacterial pathogens. In fact, a growing number of Mn transport systems have been identified as major virulence factors, including examples where loss of Mn transporters rendered organisms closely related to *S. mutans*, such as *Streptococcus pneumoniae* and *Enterococcus faecalis,* virtually avirulent in animal infection models [27, 28]. In *S. mutans*, previous characterization of pathways associated with Mn homeostasis has been restricted to the metalloregulator SloR and the ABC-type transporter SloABC [29–33]. These studies revealed that specific binding to Fe or Mn triggered SloR to function as a global transcriptional repressor, which includes repression of the *sloABC* operon [30–32]. SloABC was shown to function as a dual Fe and Mn transporter, and virulence of a *sloA* mutant strain was attenuated in a rat model of endocarditis [33].

To further our understanding of the significance of Mn homeostasis to *S. mutans* pathobiology, we first used RNA deep sequencing (RNA-Seq) to compare the transcriptome of the *S. mutans* serotype *c* strain UA159 grown in a chemically-defined medium under Mn-depleted or Mn-replete conditions. Among the genes highly upregulated during Mn starvation were all genes of the *sloABC* operon and *smu770c*, herein *mntH*, coding for a putative metal transporter from the <u>N</u>atural <u>R</u>esistance-<u>A</u>ssociated <u>M</u>acrophage <u>P</u>rotein-type (Nramp) family. While inactivation of *sloC*, coding for the SloC lipoprotein receptor, or *mntH* alone did not cause a significant impact in the overall fitness of *S. mutans*, simultaneous inactivation of *sloC* and *mntH* (Δ*sloC*Δ*mntH* strain) resulted in a dramatic reduction in cellular Mn and impaired growth and survival when cells were grown under Mn-restricted conditions. Further characterization of the Δ*sloC*, Δ*mntH,* and Δ*sloC*Δ*mntH* strains revealed that Mn transport contributes to the ability *of S. mutans* to cope with acid and oxidative stresses and to form biofilms in the presence of sucrose. Collectively, this study reveals that Mn transport in *S. mutans* is primarily mediated by SloABC and MntH and support the idea that Mn plays a critical role in the expression of virulence attributes by this important human pathogen.

## RESULTS

### Transcriptome analysis reveals a new Mn transporter in *S. mutans*

Comparison of the transcriptome profile of UA159 grown to mid-exponential phase in a chemically-defined media depleted for Mn (∼ 0.2 μM Mn) versus Mn-replete (∼ 130 μM Mn) conditions identified 95 differentially expressed genes (Table 1, False Discovery Rate (FDR) 0.01, 2-fold cutoff). Among those, 33 genes were upregulated and 62 were downregulated. To ensure that these gene expression trends were indeed due to Mn restriction, the intracellular Mn content of *S. mutans* UA159 grown under Mn-replete or Mn-depleted conditions was determined using inductively coupled plasma optical emission spectrometry (ICP-OES). The analysis confirmed that intracellular Mn content was severely diminished when *S. mutans* UA159 was grown in the Mn-depleted FMC (Fig. 1A). The residual Mn detected in cells grown in the Mn-depleted medium was likely due to carryover from the BHI medium in which overnight cultures were grown.

**FIG 1.**
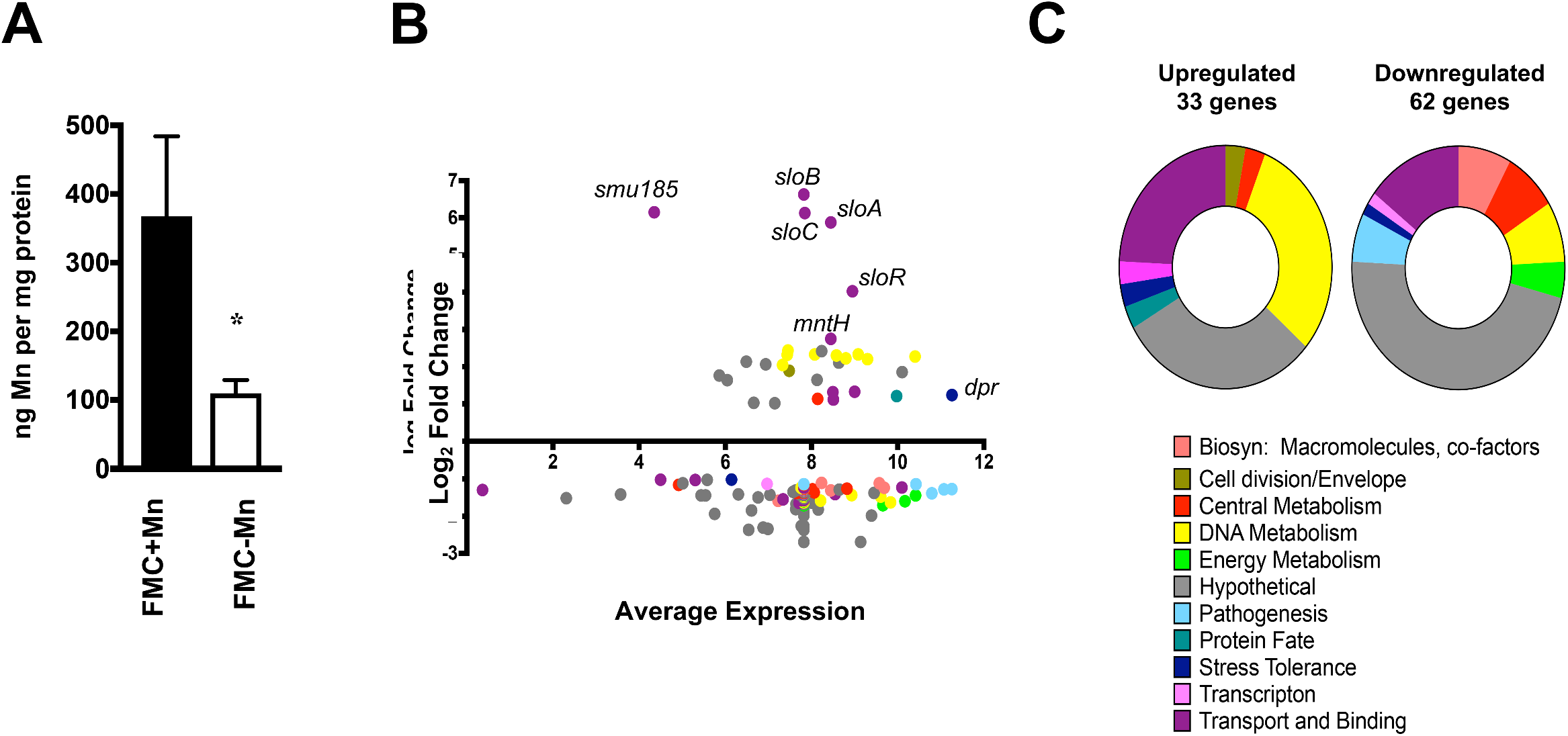
Summary of RNA-Seq analysis comparing *S. mutans* UA159 grown under Mn depleted versus Mn replete conditions. *S. mutans* UA159 was grown to an OD_600_ of 0.4 in FMC medium complete or depleted of Mn. Total RNA was isolated and gene expression under each condition was compared via RNA-Seq analysis. (A) Intracellular Mn content of *S. mutans* UA159 grown to an OD_600_ ∼0.4 in FMC medium complete or depleted of Mn. The bar graphs show the average and standard deviations of five independent ICP-OES analyses. Student’s *t*-test was used to compare metal content between the two media (*, *p* ≤ 0.005). (B) Dot plot of genes differentially expressed under conditions of Mn depletion as determined by Degust (degust.erc.monash.edu). The y-axis indicates the log_2_ fold change in expression as compared to control cultures (FMC complete), while the x-axis indicates the average expression level of each gene as compared to all other genes. The identities of selected genes of interest are indicated. (C) Graphical representations of the functional categories for up- or down-regulated genes shown in part B.

**Table 1.** *S. mutans* genes differentially expressed when grown in FMC depleted of Mn as compared to FMC complete media.

| Locus | Gene Name, Function | Fold Change | p-value |
| --- | --- | --- | --- |
| **Upregulated** |  |  |  |
| SMU_0082 | *dnaK*, chaperone protein | 2.31 | 7.03E-06 |
| SMU_0182 | *sloA*, ABC transporter, ATP-binding protein | 58.87 | 5.58E-16 |
| SMU_0183 | *sloB*, ABC transporter permease element | 99.02 | 1.34E-16 |
| SMU_0184 | *sloC*, ABC transporter, substrate binding protein | 70.07 | 1.43E-17 |
| SMU_0185 | hypothetical protein | 71.05 | 4.86E-13 |
| SMU_0186 | *sloR*, metal-dependent transcriptional regulator | 16.30 | 1.73E-16 |
| SMU_0438c | (R)-2-hydroxyglutaryl-CoA dehydratase activator-related protein | 2.20 | 2.27E-04 |
| SMU_0503c | hypothetical protein | 3.39 | 9.07E-09 |
| SMU_0540 | *dpr*, peroxide resistance protein/iron binding protein | 2.36 | 3.82E-07 |
| SMU_0600c | conserved hypothetical protein | 2.04 | 4.46E-06 |
| SMU_0609 | *bsp*, cell wall protein precursor | 3.71 | 4.34E-11 |
| SMU_0635 | conserved hypothetical protein | 4.32 | 4.98E-12 |
| SMU_0768c | conserved hypothetical protein | 4.40 | 2.97E-11 |
| SMU_0769 | conserved hypothetical protein | 2.03 | 1.68E-10 |
| SMU_0770c | *mntH*, manganese transporter periplasmic protein | 6.73 | 2.52E-13 |
| SMU_0941c | conserved hypothetical protein | 3.13 | 7.44E-06 |
| SMU_0984 | hypothetical protein | 3.12 | 1.07E-08 |
| SMU_0996 | *yclN*, ABC transporter, permease protein | 2.17 | 2.27E-03 |
| SMU_0997 | *fecE*, ABC transporter, ATP-binding protein | 2.49 | 9.29E-04 |
| SMU_0998 | *fatB*, ABC transporter, ferrichrome-binding protein | 2.51 | 5.60E-04 |
| SMU_1750c | hypothetical protein | 4.19 | 1.42E-09 |
| SMU_1752c | hypothetical protein | 3.62 | 3.21E-08 |
| SMU_1753c | CRISPR2-Cas | 5.04 | 2.32E-10 |
| SMU_1754c | CRISPR2-Cas | 5.35 | 4.14E-10 |
| SMU_1755c | CRISPR2-Cas | 4.99 | 2.58E-10 |
| SMU_1757c | CRISPR2-Cas | 5.41 | 6.85E-10 |
| SMU_1758c | CRISPR2-Cas | 4.94 | 1.15E-10 |
| SMU_1760c | CRISPR2-Cas | 5.03 | 2.32E-10 |
| SMU_1761c | CRISPR2-Cas | 4.61 | 2.60E-10 |
| SMU_1762c | CRISPR2-Cas | 4.13 | 4.47E-10 |
| SMU_1763c | CRISPR2-Cas | 4.68 | 6.72E-10 |
| SMU_1764c | CRISPR2-Cas | 4.84 | 2.33E-10 |
| SMU_2027 | transcriptional regulator/repressor | 2.17 | 9.31E-08 |
| **Downregulated** |  |  |  |
| SMU_0029 | *purC*, phosphoribosylaminoimidazole-succinocarboxamide synthase | -3.00 | 3.37E-08 |
| SMU_0030 | *purL*, phosphoribosylformylglycinamidine synthase | -2.16 | 5.98E-06 |
| SMU_0191c | phage-related integrase | -2.35 | 1.35E-05 |
| SMU_0193c | conserved hypothetical protein | -2.71 | 1.92E-05 |
| SMU_0194c | conserved hypothetical protein, phage-related | -2.71 | 1.39E-06 |
| SMU_0195c | hypothetical protein | -2.66 | 2.53E-05 |
| SMU_0196c | immunogenic secreted protein (transfer protein) | -2.44 | 2.55E-05 |
| SMU_0197c | hypothetical protein | -2.59 | 2.44E-05 |
| SMU_0198c | conjugative transposon protein | -2.78 | 2.31E-05 |
| SMU_0199c | hypothetical protein | -2.81 | 1.09E-05 |
| SMU_0200c | hypothetical protein | -2.68 | 3.57E-05 |
| SMU_0201c | conserved hypothetical protein | -2.99 | 6.44E-06 |
| SMU_0202c | conserved hypothetical protein | -3.11 | 1.39E-06 |
| SMU_0204c | hypothetical protein | -3.58 | 2.68E-06 |
| SMU_0205c | conserved hypothetical protein | -3.95 | 2.06E-07 |
| SMU_0206c | hypothetical protein | -2.48 | 4.70E-05 |
| SMU_0207c | transcriptional regulator | -2.70 | 2.90E-05 |
| SMU_0208c | conserved hypothetical protein, FtsK/SpoIIIE family | -3.09 | 9.09E-06 |
| SMU_0209c | hypothetical protein | -3.18 | 8.74E-07 |
| SMU_0210c | hypothetical protein | -2.66 | 2.61E-05 |
| SMU_0211c | hypothetical protein | -3.36 | 2.07E-05 |
| SMU_0212c | hypothetical protein | -3.83 | 6.20E-06 |
| SMU_0213c | hypothetical protein | -5.15 | 1.30E-06 |
| SMU_0214c | hypothetical protein | -4.93 | 2.04E-06 |
| SMU_0215c | hypothetical protein | -5.06 | 6.35E-07 |
| SMU_0216c | hypothetical protein | -4.80 | 2.06E-06 |
| SMU_0217c | conserved hypothetical protein | -6.46 | 2.22E-07 |
| SMU_0218 | transcriptional regulator | -2.19 | 5.25E-08 |
| SMU_0651c | ABC transporter, substrate-binding protein | -2.03 | 3.51E-03 |
| SMU_0653c | *tauC*, ABC transporter, permease protein | -2.03 | 8.83E-04 |
| SMU_0910 | *gtfD*, glucosyltransferase-S | -2.71 | 4.52E-11 |
| SMU_0932 | conserved hypothetical protein | -3.50 | 1.38E-04 |
| SMU_0933 | *atmA*, amino acid substrate-binding protein | -3.12 | 4.40E-04 |
| SMU_0934 | amino acid ABC transporter, permease protein | -2.96 | 8.67E-04 |
| SMU_0935 | amino acid ABC transporter, permease protein | -2.92 | 7.30E-04 |
| SMU_0936 | amino acid ABC transporter, ATP-binding protein | -2.87 | 6.37E-04 |
| SMU_0961 | macrophage infectivity potentiator-related protein | -3.52 | 2.72E-07 |
| SMU_0962 | *mmgC*, acyl-CoA dehydrogenase | -3.26 | 1.37E-06 |
| SMU_0992 | hypothetical protein | -2.53 | 8.40E-10 |
| SMU_1072c | *bar*, acyltransferase | -2.23 | 3.18E-07 |
| SMU_1284c | conserved hypothetical protein | -2.03 | 4.15E-08 |
| SMU_1286c | *blt*, multidrug resistance permease | -2.02 | 2.76E-08 |
| SMU_1334 | *mubP*, phosphopantetheinyl transferase | -2.42 | 4.37E-09 |
| SMU_1335c | *mubJ*, enoyl-acyl carrier protein reductase | -2.39 | 3.57E-10 |
| SMU_1336 | *mubI*, conserved hypothetical protein | -2.56 | 2.38E-09 |
| SMU_1337c | *mubM*, alpha/beta superfamily hydrolases | -2.59 | 1.76E-10 |
| SMU_1338c | *mubZ*, ABC transport macrolide permease | -2.65 | 8.92E-09 |
| SMU_1339 | *mubD*, bacitracin synthetase | -2.61 | 3.28E-09 |
| SMU_1340 | *mubC*, bacitracin synthetase 1 | -2.42 | 3.12E-08 |
| SMU_1341c | *mubB*, gramicidin S synthase | -2.20 | 9.82E-08 |
| SMU_1342 | *mubA*, bacitracin synthetase | -2.41 | 2.80E-08 |
| SMU_1343c | *mubH*, polyketide synthase | -2.34 | 3.92E-07 |
| SMU_1344c | *fabD*, malonyl CoA-acyl carrier protein transacylase | -2.47 | 9.22E-07 |
| SMU_1345c | *mycA*, peptide synthetase | -2.35 | 1.57E-06 |
| SMU_1346 | *mubT*, thioesterase II-like protein | -2.15 | 1.21E-05 |
| SMU_1395c | hypothetical protein | -2.85 | 2.01E-06 |
| SMU_1895c | hypothetical protein | -2.51 | 4.50E-07 |
| SMU_1896c | hypothetical protein | -2.72 | 9.12E-09 |
| SMU_1899 | ABC transport fragment | -2.45 | 1.89E-03 |
| SMU_1912c | hypothetical protein | -2.16 | 3.89E-04 |
| SMU_2028 | *ftf*, fructosyltransferase | -3.02 | 2.97E-09 |
| SMU_2076c | hypothetical protein | -2.67 | 1.63E-07 |

The differentially expressed genes were grouped into 11 functional categories (Fig. 1 B and C), with genes encoding transport and binding, DNA metabolism, and hypothetical proteins highly represented in the list of upregulated genes. In contrast, genes encoding hypothetical proteins accounted for more than 50% of the downregulated genes followed by genes involved in transport and binding. The most highly upregulated genes during growth under Mn-restricted conditions were those of the dual Fe and Mn transporter *sloABC* operon (≥56 to 99-fold), a small open reading frame (*smu185*, 71-fold) with the first 18 nucleotides overlapping the *sloC* gene 3’ end, the *sloR* transcriptional repressor (16-fold), and the uncharacterized *smu770c* (6-fold) (Table 1, Fig. 1B). BLAST search analysis revealed that the protein encoded by *smu770c* belongs to the Nramp-type transport family predicted to function in metal uptake. The Smu770c protein shared 76% identity with the *S. agalactiae* (Group B *Streptococcus*) MntH and 60 and 54% identity with *E. faecalis* MntH1 and MntH2 proteins, respectively. Of note, the *S. agalactiae* MntH and *E. faecalis* MntH1 and MntH2 have been recently assigned a role in Mn uptake [27, 34]. Other genes upregulated in the absence of Mn were several belonging to the CRISPR2-*cas* operon (*smu1753c*-*smu1764c*; >4-fold), as well as 3 of 4 genes from the *smu995-998* operon (>2-fold), recently shown to code for an Fe transport system [35].

The most repressed genes when *S. mutans* was grown under Mn restricted conditions were a cluster of genes encoding possible conjugative transposon proteins (*smu191c*-*217c*, ≥2.4-fold downregulated). Additionally, genes encoding proteins with predicted roles in amino acid transport (*smu932-936*), purine biosynthesis (*smu29-smu32*), fatty acid biosynthesis (*smu1334c-smu1338c*), production of antimicrobial compounds (*smu1339c-1343c*), and sugar transport and metabolism (*ftf*, *smu2028*, *gtfD*, *smu910*) showed decreased levels of expression under Mn-depleted conditions (Table 1).

### SloABC and MntH are the principal manganese transporters in *S. mutans*

Because of the high degree of conservation between Smu770c and previously characterized MntH proteins from other Firmicutes, we assigned the name *“mntH”* to the monocistronic transcriptional unit *smu770c*. Here, we sought to characterize the *mntH* gene and investigate the possible cooperative nature of SloABC and MntH in metal acquisition. To accomplish this, we created strains bearing single deletions in *sloC (*Δ*sloC*), which encodes the metal binding lipoprotein of the SloABC system, or in *mntH* (Δ*mntH*), as well as a double mutant strain lacking both *sloC* and *mntH* (Δ*sloC*Δ*mntH*). Because BHI contains fairly low levels of Mn (∼0.6 μM) (Table 2) and, in anticipation that some of the mutant strains would grow poorly or not grow in plain BHI, all mutant strains were initially isolated on BHI agar supplemented with 75 μM Mn. Upon genetic confirmation of the single and double mutants, we tested the ability of these strains to grow in BHI and found that the Δ*sloC*Δ*mntH* double mutant was unable to grow on BHI agar without Mn supplementation (Fig. 2A). The Δ*sloC*Δ*mntH* strain was able to grow in BHI broth, albeit at much slower rates when compared to the other strains, reaching similar final growth yields after 16 hours (Fig. 2B). Supplementation of BHI with 25 μM Mn (BHI+Mn) fully restored the growth defect of the double mutant strain in broth (Fig. 2C). We suspected that the different growth behavior of the Δ*sloC*Δ*mntH* strain in BHI plates or broth was due to trace amounts of Mn that transferred from the overnight BHI inoculum that contained 7 μM Mn. This suspicion was then confirmed by findings that the Δ*sloC*Δ*mntH* strain could not grow in unsupplemented BHI after a second passage (data not shown). To assess the metal requirements of the mutant strains in a more controlled fashion, growth of the parent UA159 and mutant strains was also monitored in the chemically-defined FMC medium (Fe- and Mn-replete, Table 2), and in FMC depleted of Mn (Mn < 90 nM), Fe (Fe < 90 nM), or both [27]. In complete FMC, growth of all mutant strains was indistinguishable from that of the parent strain (Fig. 2D). As expected, the Δ*sloC*Δ*mntH* double mutant strain failed to grow in Mn-depleted FMC whereas the Δ*sloC* mutant showed a slight growth delay that did not affect final growth yields (Fig. 2E). Iron depletion alone did not affect growth of the parent or any of the mutant strains, but simultaneous depletion of Fe and Mn exacerbated the slow growth defect of the Δ*sloC* strain (Fig. 2F-G). Growth of the Δ*sloC*Δ*mntH* strain in plain BHI or in Mn-depleted FMC was fully restored by complementation, when either the *sloC* or *mntH* gene was integrated elsewhere in the chromosome (Fig. 2A and H).

**FIG 2.**
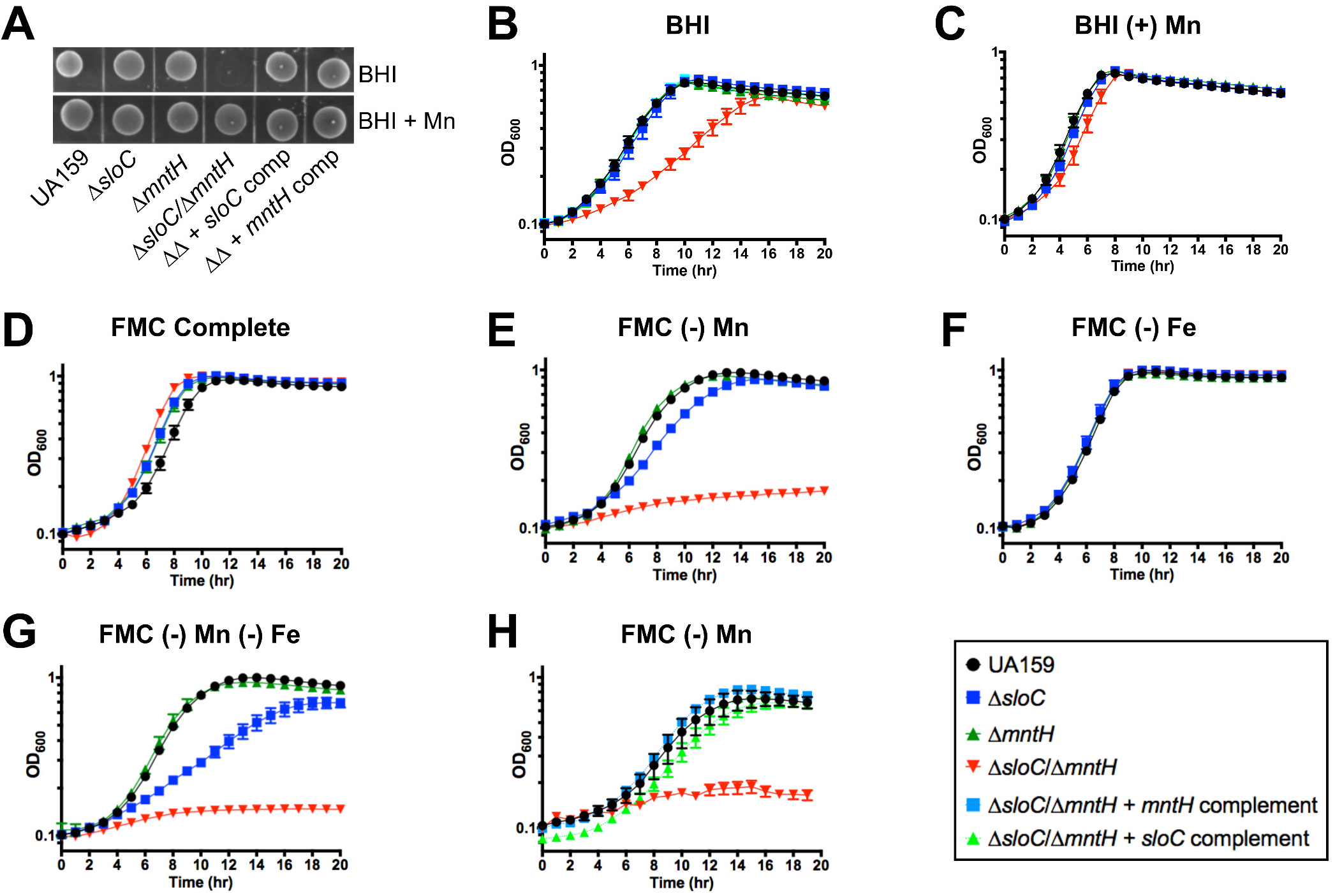
SloABC and MntH promote growth of *S. mutans* in Mn-depleted environments. (A) Growth of *S. mutans* UA159, Δ*sloC*, Δ*mntH,* and *Δsloc/ΔmntH* mutant strains along with the double mutant strain complemented with either *sloC* or *mntH* to mid-logarithmic phase (OD_600_ ∼0.4) on BHI agar. Overnight cultures were spotted onto BHI agar with or without supplementation with 10 μM Mn. Plates were incubated for 48 hours before image was obtained. (B-G) Growth of UA159, Δ*sloC*, Δ*mntH,* and *Δsloc/ΔmntH* mutant strains in (B) BHI broth, (C) BHI broth supplemented with 75 μM Mn, (D) FMC complete (130 μM Mn), (E) Mn-depleted FMC, (F) Fe-depleted FMC, and (G) Mn- and Fe-depleted FMC. (H) Genetic complementation of the *ΔsloC/ΔmntH* growth defect in Mn-depleted FMC with either *sloC* or *mntH.* The graphs show the average and standard deviations of at least three independent experiments.

**Table 2.** Metal content of media used for growth of *S. mutans*.

| Metal | BHI | FMC | Saliva |
| --- | --- | --- | --- |
| Iron | 5.91 ± 1.27 µM | 82.62 ± 8.8 µM | 4.51 ± 0.08 µM |
| Manganese | 0.56 ± 0.27 µM | 132.6 ± 14.9 µM | BDL* |
| Zinc | 10.9 ± 2.01 µM | 1.2 ± 0.3 µM | 0.4 ± 0.02 µM |

ICP-OES analysis was used to determine the metal content of BHI, FMC and pooled human saliva used in this study. Values represent the averages and standard deviations of at least three independent experiments.\* BDL: below detection limit

Next, we used ICP-OES to determine the cellular metal content of parent and mutant strains grown to mid-exponential phase in BHI broth (Fig. 3). Despite not showing a growth defect in plain BHI, the Δ*sloC* and Δ*mntH* single mutant strains carried ∼ 45% less cellular Mn when compared to UA159. In agreement with the results shown in Figure 2, combined deletion of *sloC* and *mntH* resulted in a more significant reduction (∼ 80% less) in cellular Mn pools. Complementation of Δ*sloC*Δ*mntH* with either one of the inactivated genes restored cellular Mn content to parent strain levels. Despite the previously assigned role of SloABC in Fe uptake [33], intracellular quantities of Fe did not vary significantly among the strains. Collectively, these results reveal that SloABC and MntH comprise the principal Mn transport systems of *S. mutans,* working cooperatively to maintain Mn homeostasis.

**FIG 3.**
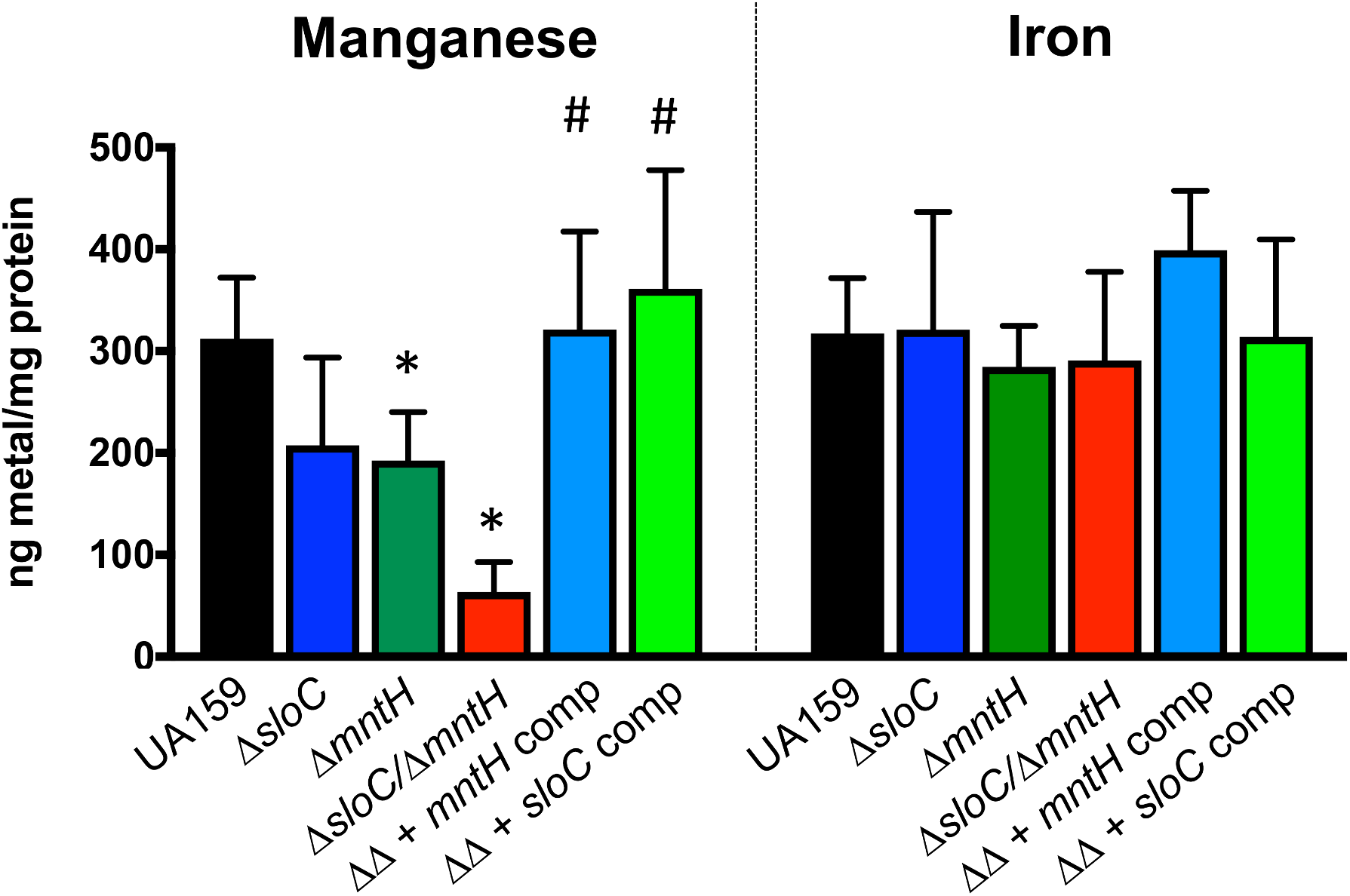
SloABC and MntH are the main Mn transporters in *S. mutans* UA159. Intracellular Mn and Fe content of *S. mutans* UA159 and derivatives grown in plain BHI to an OD_600_ ∼ 0.4. The bar graphs show the average and standard deviations of five independent ICP-OES analyses. Student‘s *t*-test was used to compare metal content of the mutant strains to that of UA159 (*, *p* ≤ 0.05), and the double mutant *ΔsloCΔmntH* to the complemented strains (#, *p* ≤ 0.0005).

### *mntH* is a new member of the SloR regulon

Transcriptional repression of the *sloABC* operon exerted by SloR has been thoroughly characterized by one of our laboratories [30, 31, 36]. In one of these studies a conserved SloR-binding palindrome was identified upstream of the *mntH* gene but the specificity of SloR binding to the *mntH* promoter region was not explored at that time [31]. Here, we used quantitative real time PCR (qRT-PCR) and an electrophoretic mobility shift assay (EMSA) to determine the SloR-*mntH* relationship. When compared to the parent strain, inactivation of *sloR* (Δ*sloR* strain) increased *mntH* transcription by ∼ 5-fold and of *sloA,* the first gene of the *sloABC* operon, by ∼ 15-fold (Fig. 4A). In addition, EMSAs revealed that as little as 60 nM of purified SloR shifted *mntH* probe migration (Fig. 4B), and that the region possibly harbors more than a single SloR binding site given the supershift that was observed with 300 nM SloR. The specificity of SloR binding to the *mntH* probe was confirmed by showing that addition of the metal chelator EDTA, or excess cold probe disrupts the interaction in a concentration-dependent manner (Fig. 4B).

**FIG 4.**
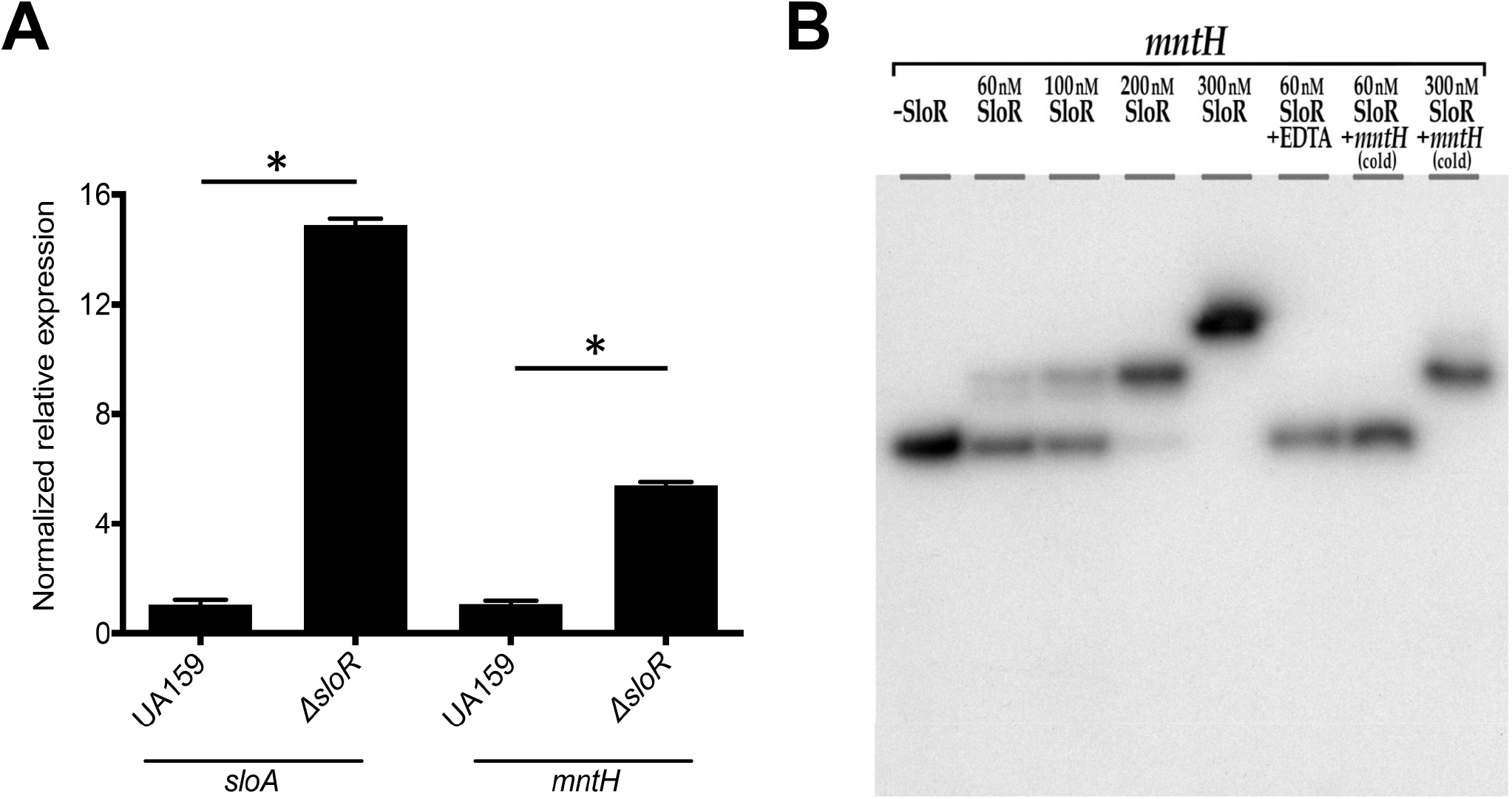
The *S. mutans mntH* gene belongs to the SloR regulon. (A) qRT-PCR analysis indicate expression levels of *mntH* and *sloA* that are up-regulated in a *ΔsloR* strain when compared to the parent strain UA159. Shown are the means ± and standard deviations of 3 independent experiments. Student‘s *t*-test was used to compare differences in gene expression between UA159 and *ΔsloR* strains. (B) Regulation of the *S. mutans mntH* gene by SloR is direct. EMSA was performed with a [γ-^32^P] ATP end-labeled *mntH* probe and purified SloR. Reaction mixtures were resolved on 12% non-denaturing polyacrylamide gels and exposed to X-ray film for 24 h at −80°C. The addition of cold competitor DNA (1:1) or 3 mM EDTA in the SloR-*mntH* reaction mixture abrogated the band shift, whereas addition of 300 nM SloR resulted in a supershift.

### Manganese is critical for *S. mutans* tolerance to clinically relevant conditions

To examine the importance of Mn in the oxidative stress tolerance of *S. mutans*, we first grew cells in the presence of a sub-inhibitory concentration of H_2_O_2_. Under the conditions tested, growth of parent, Δ*sloC* or Δ*mntH* strains was not affected; however, growth rates and yields of the Δ*sloC*Δ*mntH* double mutant strain were markedly reduced (Fig. 5A). Importantly, this growth defect was rescued by Mn supplementation (Fig. 5B). In parallel, we tested this same panel of strains in a qualitative competition assay against the net H_2_O_2_ producing oral commensals *Streptococcus gordonii* and *Streptococcus sanguinis*. While the antagonizing peroxigenic strain inhibited growth of all strains, the growth inhibition of the Δ*sloC*Δ*mntH* strain was much more pronounced (Fig. 5C). The inhibitory effect of the peroxigenic streptococci was abolished by the addition of of catalase.

**FIG 5.**
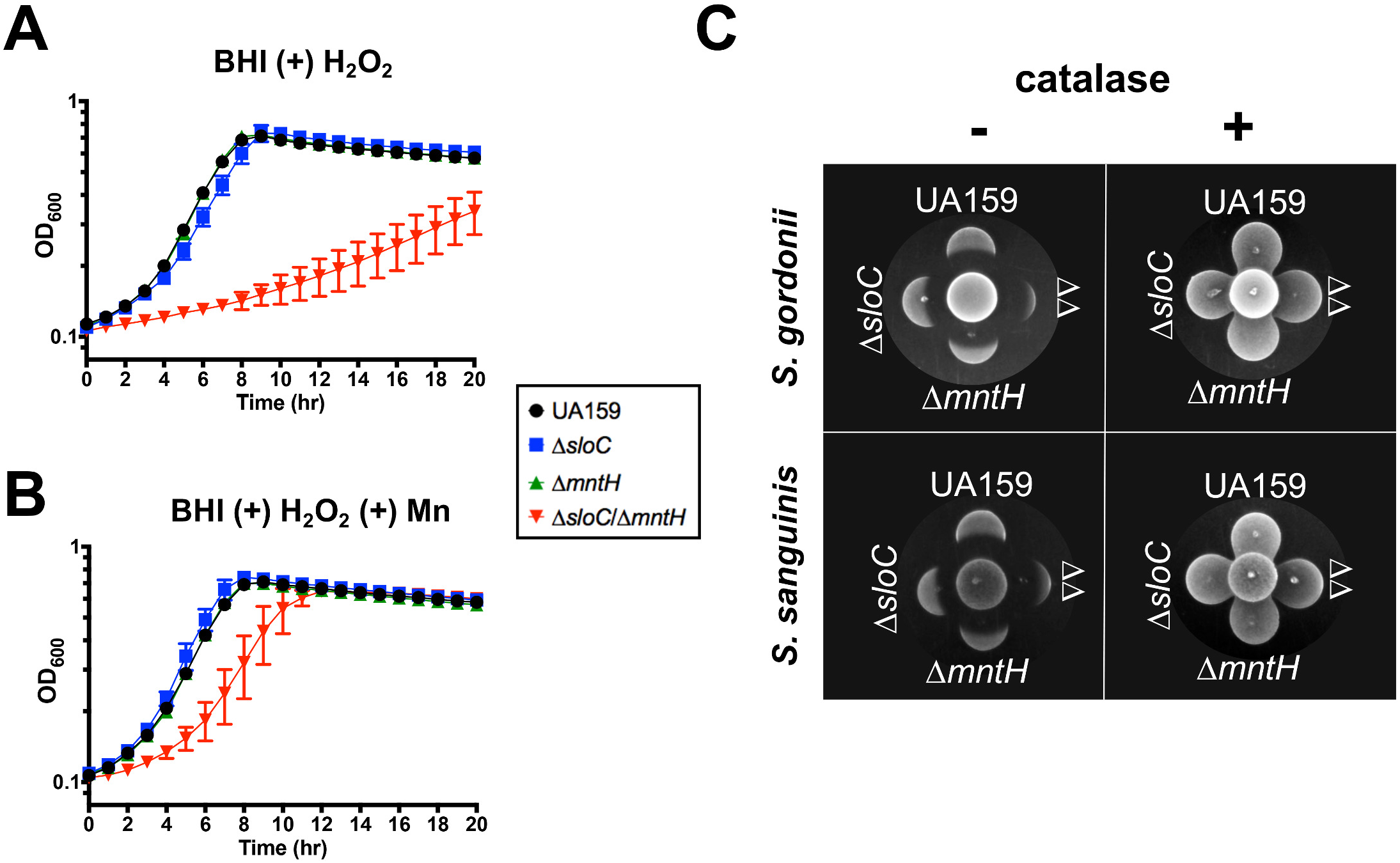
Manganese transport contributes to H_2_O_2_ tolerance. (A) Growth of *S. mutans* UA159, Δ*sloC*, Δ*mntH* and *ΔslocΔmntH* strains in the presence of 0.2 mM H_2_O_2_ in (A) plain BHI, or (B) BHI supplemented with 10 μM Mn. (C) Peroxigenic strains *S. gordonii* DL-1 or *S. sanguinis* SK150 were spotted at the center of a BHI agar plate (supplemented with 2 µM Mn) and grown for 24 hours (37°C, 5% CO_2_). *S. mutans* cultures were then spotted proximal to the peroxigenic strain and grown for an additional 24 hours. The center spot of each grouping shown here is the H_2_O_2_-producing strain, while the *S. mutans* strains are labeled in the figure (*ΔΔ* corresponds to the Δ*sloC*Δ*mntH* double mutant). As a control, duplicate spotting was performed in which H_2_O_2_ produced by the peroxigenic strains was neutralized by overlaying the innoculum spot with a catalase solution prior to spotting *S. mutans*. The images shown are representative of three independent experiments.

The ability to withstand acid stress is a major virulence attribute of *S. mutans* that sets it apart as a cariogenic organism when compared to less aciduric commensal streptococci. Recently, the *S. agalactiae* MntH was shown to play a crucial role in low pH survival [34]. To probe the significance of Mn in acid stress, cultures of parent and mutant strains were grown in FMC adjusted to pH 7.0 (control) or pH 5.5 (acid stress) containing the concentration of Mn indicated in the original recipe (130 μM Mn), or with the minimal concentration of Mn (3 μM Mn) that sustains optimal growth of the Δ*sloC*Δ*mntH* strain in FMC (Fig. 6A). In medium adjusted to pH 7.0, all strains grew well and reached the same final growth yield under high or low Mn conditions (data not shown). In medium adjusted to pH 5.5, all strains reached similar final growth yields in the high Mn condition (130 μM Mn) (Fig. 6B). However, all strains showed reduced final growth yields in the low Mn medium adjusted to pH 5.5 (when compared to high Mn medium). Moreover, the final growth yields of the Δ*mntH* and Δ*sloC*Δ*mntH* strains was further impaired in the low Mn medium adjusted to pH 5.5 (Fig. 6B). Collectively, these results reveal that a minimal threshold of intracellular Mn is determining for the oxidative and acid stress tolerance of *S. mutans*.

**FIG 6.**
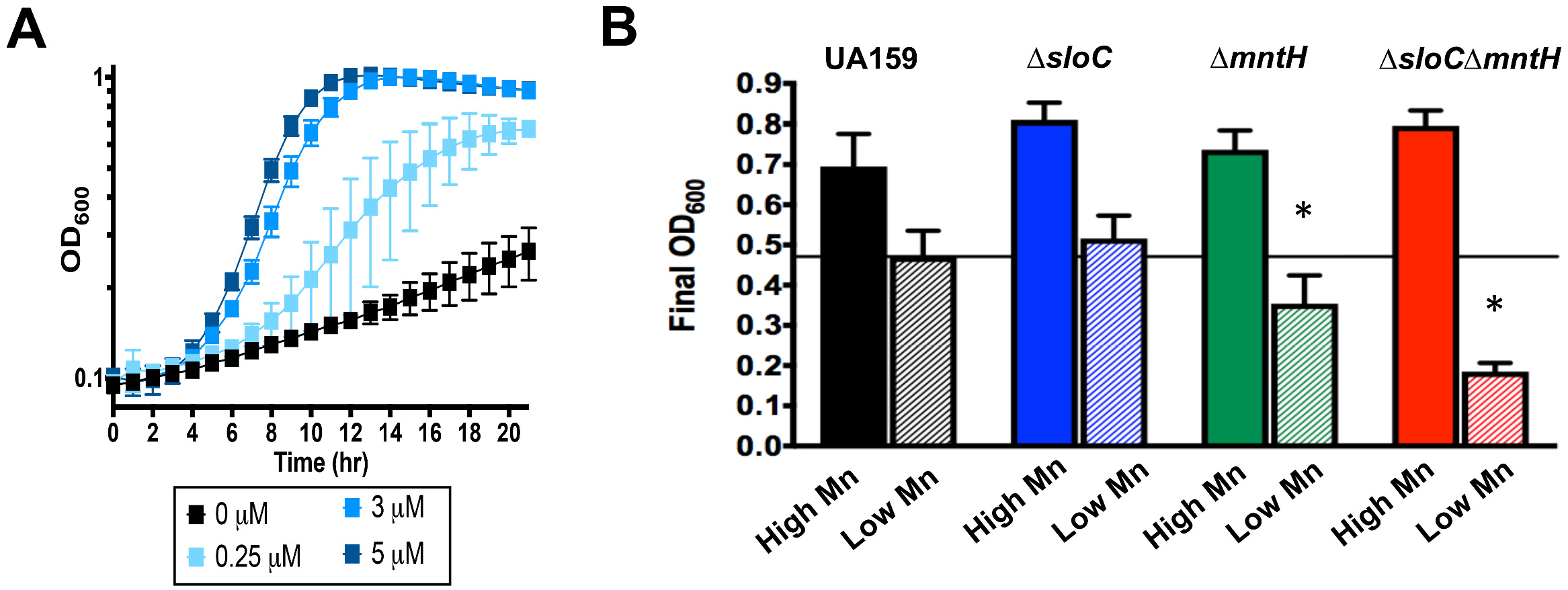
Manganese transport contributes to acid stress tolerance in *S. mutans*. (A) Growth curves showing the minimal concentration of Mn that fully supports growth of the Δ*sloC*/Δ*mntH* strain. The graphs represent the average and standard deviations of three independent cultures. (B) Growth of *S. mutans* UA159, Δ*sloC*, Δ*mntH*, or Δ*sloC*Δ*mntH* in FMC medium adjusted to pH 5.5 containing ∼ 130 µM Mn (High Mn; solid bars), or 3 µM Mn (low Mn; striped bars). Bars represent the mean and standard deviation of the final OD_600_ values for five independent experiments. The horizontal line represents the mean final OD_600_ for UA159 grown in FMC containing low Mn. Student‘s *t*-test was used to compare the final values of the mutant strains to that of UA159 grown in the same medium. (*) *p* < 0.05.

### Manganese promotes sucrose-dependent biofilm formation

Next, we investigated the ability of the Mn transport mutant strains to adhere and form biofilms on saliva-coated microtiter plate wells using BHI supplemented with 2% sucrose. In the early stage of biofilm development (4 h incubation), all mutant strains showed a significant defect in biofilm formation, with the Δ*sloC*Δ*mntH* strain showing the most pronounced defect (∼ 85% reduction) (Fig. 7A). Supplementation of the growth media with Mn partially restored the early stage biofilm defect of the double mutant strain (Fig. 7A). After the mature biofilm was formed (24 h incubation), only the Δ*sloC*Δ*mntH* double mutant continued to show a statistically significant defect in biofilm formation (∼ 25% reduction); this phenotype was fully restored by Mn supplementation (Fig. 7B). Despite the slow growth phenotype of the Δ*sloC*Δ*mntH* double mutant in BHI broth (Fig. 2B), no differences in growth (based on OD_600_ and CFU counts) were observed among strains in the two time points shown in Figure 7 (data not shown). Collectively, these results support previous observations indicating that the ability to maintain intracellular Mn homeostasis is important for sucrose-dependent biofilm formation.

**FIG 7.**
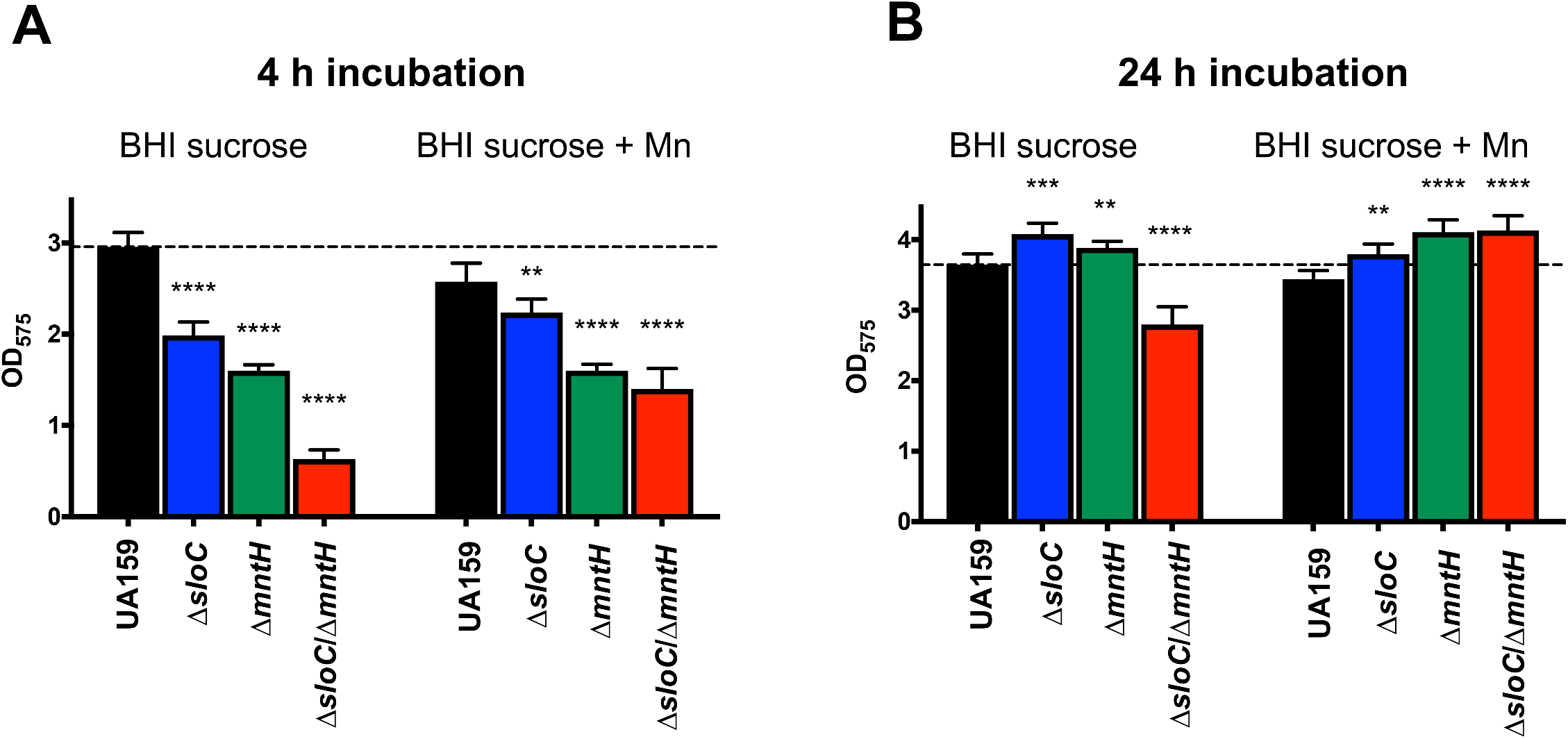
Manganese acquisition is important for sucrose-dependent biofilm formation of *S. mutans* UA159. Cultures were gown in BHI broth containing 2% sucrose with or without supplementation with 10 μM Mn for 4- or 24-h in saliva-coated microtiter wells. The graph shows the average and standard deviation of three independent experiments performed in quadruplicate. (**) *p* ≤ 0.05, (***) *p* ≤ 0.01, (****) *p* ≤ 0.005.

### Growth and survival of the Δ*sloC*Δ*mntH* strain was impaired in human saliva *ex vivo*

As a resident of the human oral cavity, *S. mutans* is bathed in saliva; therefore, the ability to proliferate and survive in this biological fluid is an important aspect of its lifestyle. Here, we tested the ability of parent and mutant strains to grow and survive in pooled human saliva supplemented with 10 mM glucose to promote a more robust cell growth. Metal quantifications revealed that our batch of pooled saliva had relatively high Fe (4.51 ± 0.08 μM) Fe and low Zn (0.4 ± 0.02 μM) levels while the level of Mn was below the detection limit (Table 2). The parent and single mutant strains grew well in saliva, showing a peak increase in CFU of nearly 2 logs of growth within the initial 18 h, followed by a noticeable loss of cell viability after 48 h (Fig. 8A). On the other hand, the Δ*sloC*Δ*mntH* strain grew poorly within the initial few hours and rapidly lost viability, eventually yielding no viable cells by 48 h. Supplementation of the saliva-glucose media with 10 μM Mn allowed all strains (including Δ*sloC*Δ*mntH*) to reach maximal growth yields faster and maintain viability comparable to that of the parent strain during the initial 24 h (Fig. 8B).

**FIG 8.**
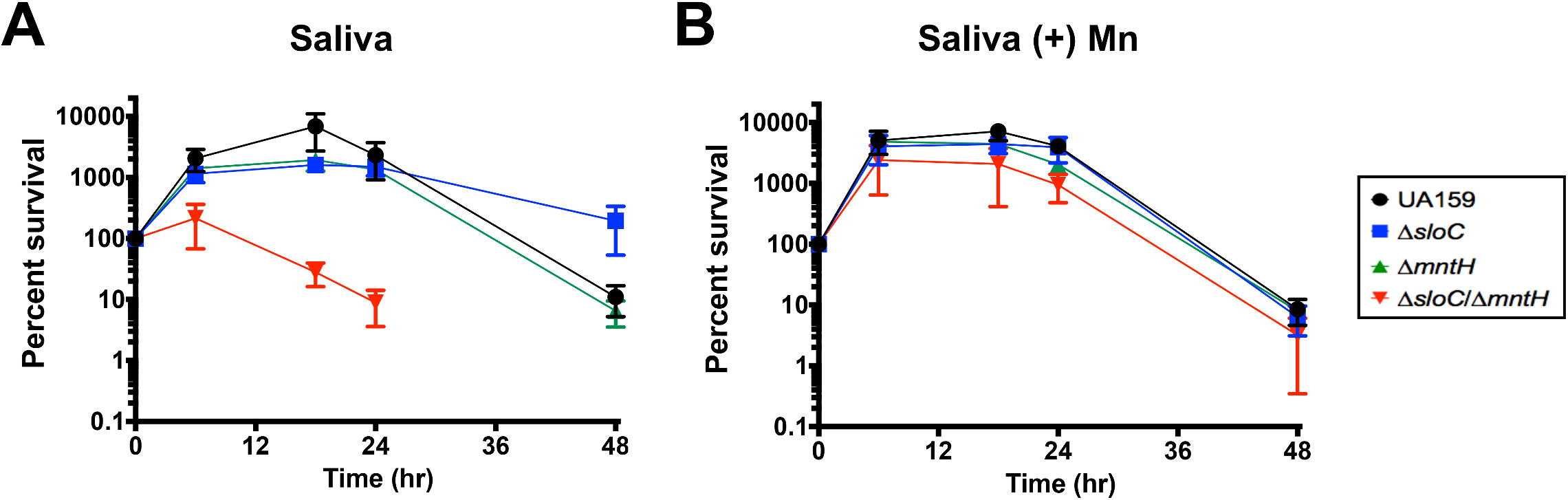
Manganese transport is critical for *S. mutans* growth and survival in human saliva. Strains (UA159, Δ*sloC*, Δ*mntH* or Δ*sloC*Δm*ntH*) were grown in plain BHI to OD_600_ ∼ 0.3, washed in PBS and diluted 1:20 in (A) pooled saliva containing 10 mM glucose, or (B) pooled saliva supplemented with 10 mM glucose and 10 μM Mn. The graphs show the average and standard deviation of four independent experiments.

### SloABC and MntH are required for calprotectin tolerance

The bioavailability of metals in body fluids is largely dependent on the presence and activity of metal-sequestering proteins such as transferrin, lactoferrin and calprotectin. In the case of Mn, calprotectin is the major host protein responsible for sequestering Mn (as well as Zn) during infection [3]. Normally found in circulating blood and tissues at low levels, calprotectin accumulates to concentrations of up to 1 mg ml^-1^ in response to inflammation and infection, thereby playing a central role in host-activated nutritional immunity. Here, we tested the ability of *S. mutans* parent and Mn transport mutants to grow in the presence of sub-inhibitory concentrations of purified calprotectin (Fig. 9). We found that 150 μg ml ^-1^ calprotectin significantly delayed growth of the Δ*sloC* mutant, and nearly abolished growth of the Δ*sloC*Δ*mntH* double mutant (Fig. 9B). At 200 μg ml ^-1^ of calprotectin, growth of both Δ*sloC* and Δ*sloC*Δ*mntH* strains was fully inhibited (Fig. 9C). In contrast, the parent and Δ*mntH* strains grown in the presence of calprotectin showed an extended lag phase when compared to cells grown in calprotectin-free media, that did not impact final growth yields when compared to cells grown under control conditions (Fig. 9A-C). Finally, the growth inhibitory effect of calprotectin at 200 μg ml ^-1^ on the Δ*sloC* and Δ*sloC*Δ*mntH* strains could be fully overcome by supplementation with 20 μM Mn (Fig. 9D).

**FIG 9.**
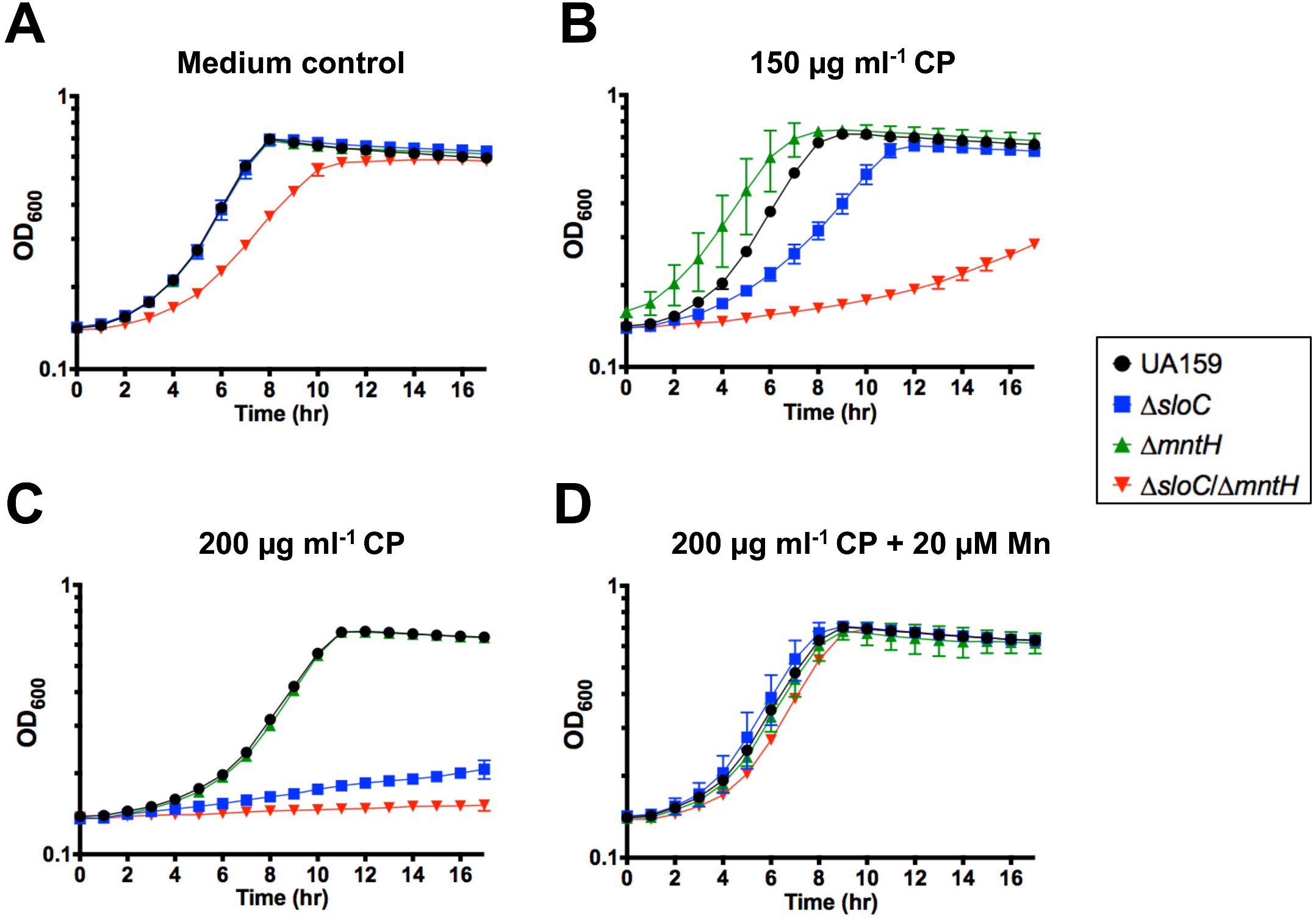
SloABC and MntH are required for *S. mutans* tolerance to calprotectin. Growth of UA159 and its derivatives in the presence of purified calprotectin (CP). Overnight cultures were diluted 1:20 into BHI, grown to early log phase (OD_600_ = 0.25), then diluted 1:50 in CP medium containing (A) no CP, (B) 150 µg ml^-1^ CP, (C) 200 µg ml^-1^ CP, or (D) 200 µg ml^-1^ CP + 20 μM Mn. The graphs show the average and standard deviations of three independent cultures.

## DISCUSSION

In this study, we showed that Mn is an essential micronutrient for *S. mutans* and that the ability to maintain Mn homeostasis is important for the expression of virulence factors associated with oral and non-oral infections. Global transcriptional profiling of *S. mutans* UA159 grown under Mn-depleted conditions led to the identification of a previously uncharacterized Mn transporter, here named MntH, belonging to the Nramp family of transporters. By studying the physiology of the Δ*sloC*, Δ*mntH* and Δ*sloC*Δ*mntH* strains, we provided unequivocal evidence that SloABC and MntH are the primary Mn transporters in *S. mutans*, and that simultaneous inactivation of *sloC* and *mntH* impaired the fitness of *S. mutans* under Mn-restricted conditions. However, the Δ*sloC*Δ*mntH* double mutant strain retained the ability to grow under Mn-rich conditions suggesting that there are alternative transporter(s), perhaps bearing high affinity for metals other than Mn, that can take up Mn when provided in excess.

Although Nramp-type proteins have been shown to transport different types of trace metal ions such as Fe, Mn and Zn [37], recent studies with *S. agalactiae* and *E. faecalis* revealed that the closest homologs of the *S. mutans* MntH are primarily involved in Mn transport [27, 34]. This appears to be the case for the *S. mutans* MntH as the intracellular levels of Fe or Zn were not affected by *mntH* inactivation (Fig. 3 and data not shown). It is interesting to note that while Nramp transporters are commonly found in bacteria, members of this family are absent in some major pathogenic streptococci such as *S. pyogenes* and *S. pneumoniae*. On the other hand, all streptococcal genomes encode one copy of an ABC-type Mn transporter homologous to SloABC. In *S. mutans*, inactivation of *sloABC* resulted in attenuated virulence in a rat model of infectious endocarditis [33] whereas inactivation of the lone Mn transporter in *S. pneumoniae* abrogated virulence in systemic, respiratory tract and otitis media infections [28]. In *E. faecalis* OG1RF, which encodes one ABC-type (EfaCBA) and two Nramp-type (MntH1 and MntH2) Mn transporters, inactivation of *efaCBA* and *mntH2* virtually abolished the virulence of *E. faecalis* in mammalian models [27]. In the future, it will be useful to test the virulence potential of the *S. mutans* Δ*mntH* and Δ*sloC*Δ*mntH* strains in an animal model of infective endocarditis, as we suspect that simultaneous disruption of *mntH* and the *sloABC* operon will abrogate the ability of *S. mutans* to cause systemic infections, yielding a much more robust phenotype than the single Δ*sloABC* mutant strain displayed [33].

After *sloABC*, *mntH* and the *sloR* repressor, the next group of overexpressed genes in cells starved for Mn belonged to the CRISPR2 system (∼5-fold average gene upregulation), which is thought to provide sequence-based immunity against “invasion” by mobile genetic elements [38]. CRISPRs are often associated with a set of *cas* genes that encode proteins that mediate the defense process. In *S. mutans* UA159, deletion of the *cas* genes associated with CRISPR2 increased cell sensitivity to heat shock without affecting cell sensitivity to the virulent phage M102 [39]. A second CRISPR system present in *S. mutans* UA159, named CRISPR1, was shown to mediate tolerance toward multiple stresses, including membrane, DNA, oxidative and heat stress [39]. While the mechanism remains to be determined, it seems that CRISPR systems are intimately associated with *S. mutans* stress responses. Among the genes downregulated under the Mn-depleted condition, 38 genes belong to the genomic islands (GI) TnSmu1 (25 genes), a 23 kb region that lies adjacent to a cluster of tRNA genes, and TnSmu2 (13 genes), the largest genomic island found in UA159 [40]. While not much is known about the biological roles of these GI in *S. mutans*, TnSmu2 is responsible for the biosynthesis of a pigment important for oxidative stress tolerance [41]. It is also noteworthy that genes belonging to CRISPR systems, and TnSmu1 and TnSmu2 are also differentially expressed in strains lacking the serine protease *clpP*, the transcriptional regulator *covR*, and *cidB* from the Cid/Lrg holin/anti-holin system [42–44]. Even though ClpP, CovR and Cid/Lrg modulate diverse biological processes, they seem to share a common role in stress tolerance and adaptation. For these reasons, studies to investigate the possible association of these mobile genetic elements with metal homeostasis should be considered in the near future.

SloR was previously shown to repress transcription of the *sloABC* operon in a Mn-dependent fashion by binding to conserved palindromes that define a so-called <u>S</u>loR recognition <u>e</u>lement (SRE) in the *sloABC* promoter region [30, 31]. As a result, growth of *S. mutans* in Mn-rich media resulted in decreased *sloABC* transcription [31, 33, 45]. Previously, a genome-wide characterization of *S. mutans* UA159 identified a putative SRE in the *mntH* promoter region [29]. Here, our results confirm that SloR contributes to the regulation of *mntH*, though the results of both RNAseq and qRT-PCR analyses indicate that SloR repression of the *sloABC* operon is tighter than it is for the *mntH* gene. Such robust *sloABC* repression by SloR can be explained by our previous characterization of cooperative, homodimeric binding between SloR and each of three hexameric repeats that overlap the *sloABC* promoter [46]. Whether SloR binding at the *mntH* locus is cooperative, and whether the SloR binding sites overlap with the *mntH* promoter remains to be determined. While the EMSA results described herein support more than a single SloR binding site upstream of the *mntH* gene, how this might translate into greater promoter accessibility to RNA polymerase, and thus more relaxed *mntH* transcription warrants further investigation.

Previous epidemiological studies have associated high trace metal availability in the oral cavity with a higher caries incidence in pre-determined populations [8–12]. In particular, Mn appears to play a prominent role in host-pathogen interactions by serving as a co-factor for bacterial enzymes involved in general metabolism, DNA replication and oxidative stress tolerance [24]. The association of Mn levels with oral streptococci physiology and cariogenicity was first examined in the late 60’s and became the subject of more intensive investigations in the mid-80’s until the early 90’s. Collectively, these studies have shown that Mn (i) is an essential co-factor for both cariogenic and non-cariogenic streptococci, (ii) plays a major role in the growth of *S. mutans* at elevated oxygen levels by serving as a co-factor of the superoxide dismutase enzyme, (iii) modulates dextran-mediated aggregation in different species of oral streptococci, and (iv) stimulates carbohydrate metabolism and IPS accumulation in *S. mutans* [13, 14, 17, 18, 20, 22, 47]. Most notably, when added to drinking water, Mn resulted in a significant increase in the total number of carious lesions as well as caries severity in germ-free WAGG rats [17]. Despite the important advances made by these studies, most were conducted prior to, or in the early days of the genomic era, when the currently available tools for molecular genetic manipulations and comparative genomics were under development. Here, taking advantage of the contemporary tools available, we confirmed some of those initial discoveries, and further expanded our understanding of how Mn influences the pathophysiology of *S. mutans*. We confirmed or showed for the first time that some of the major cariogenic traits of *S. mutans*, such as acid and oxidative stress tolerance, survival in saliva, and sucrose-dependent biofilm formation, are in fact dependent on the intracellular levels of Mn. Further, we have demonstrated that manganese transporters are critical to the ability of *S. mutans* to tolerate the host immune protein calprotectin, which pathogens encounter in the oral cavity and, particularly, in the bloodstream. These results suggest that strategies to deprive *S. mutans* of Mn hold great promise in our efforts to combat this important pathogen.

## MATERIALS AND METHODS

### Bacterial strains and growth conditions

The bacterial strains used in this study are listed in Table 3. *S. mutans* UA159 and its derivatives were routinely grown in BHI agar supplemented with 75 µM MnSO_4_ at 37°C under anaerobic conditions. For physiologic analyses, bacterial inocula were prepared from overnight cultures grown in BHI supplemented with 7 µM MnSO_4_ (BHI+Mn), sub-cultured 1:20 in plain BHI (without Mn supplementation) and grown to early logarithmic phase (OD_600_ = 0.25) at 37°C in a 5% CO_2_ atmosphere. To assess the ability of *S. mutans* strains to grow in BHI or the chemically-defined FMC medium [50], cultures prepared as indicated above were diluted 1:50 into the appropriate medium in a microtiter plate with an overlay of sterile mineral oil to minimize the deleterious effects of oxygen metabolism. Growth was monitored using the BioScreen C growth reader (Oy Growth Curves) at 37°C. Growth in the presence of calprotectin requires the use of 38% bacterial medium and 62% CP buffer (20 mM Tris pH 7.5, 100 mM NaCl, 3 mM CaCl_2_, 5 mM β–mercaptoethanol). To promote the growth of *S. mutans* in the CP medium, 3X concentrated BHI was used in combination with the CP buffer. For RNA-Seq analysis, three replicate cultures of UA159 were grown overnight as described above in plain BHI medium and then sub-cultured 1:20 in complete FMC (containing 130 µM Mn) as a control, or in Mn-depleted FMC medium from which Mn was omitted from the recipe. Cultures were grown to OD_600_ of 0.4, harvested by centrifugation, and the bacterial pellets resuspended in 1 ml RNA Protect Bacterial Reagent (Qiagen). Following another centrifugation cycle, the supernatants were discarded and the pellets stored at −80°C until use.

**Table 3.** Bacterial strains used in this study.

| Strains | Relevant genotype | Source |
| --- | --- | --- |
| <u>*S. mutans*</u> |  |  |
| UA159 | parent, serotype *c* | Laboratory stock |
| UAΔ*sloC* | *smu184::Spec* | This study |
| UAΔ*mntH* | *smu770c::Erm* | This study |
| UAΔ*sloC*/Δ*mntH* | *smu184::Spec*, *smu.770c::Erm* | This study |
| GMS584 (Δ*sloR*) | *smu186::Erm* | [32] |
| Δ*sloC*/Δ*mntH*+*sloC* | *sloC* complementation of Δ*sloC*/Δ*mntH* | This study |
| Δ*sloC*Δ*mntH*+*mntH* | *mntH* complementation of Δ*sloC*/Δ*mntH* | This study |
| <u>*S. gordonii*</u> |  |  |
| DL-1 | Wild-type | Laboratory stock |
| <u>*S. sanguinis*</u> |  |  |
| SK150 | Wild-type | Laboratory stock |
| <u>*E. coli*</u> |  |  |
| DH10B | Cloning host | Laboratory stock |

### Construction of mutant and complemented strains

*S. mutans* strains lacking the *sloC* or *mntH* gene, or both, were constructed using a PCR ligation mutagenesis approach [51]. Briefly, PCR fragments flanking the region to be deleted were ligated to an antibiotic resistance cassette (erythromycin for Δ*sloC* and spectinomycin for Δ*mntH*) and the ligation mixture was used to transform *S. mutans* UA159 according to an established protocol [51]. The double mutant strain was obtained by amplifying the Δ*mntH* region and using the resulting DNA amplicon to transform the Δ*sloC* single mutant strain. Mutant strains were isolated on BHI plates supplemented with 75 µM Mn and the appropriate antibiotic(s). Gene deletions were confirmed by sequencing amplicons containing the antibiotic cassette insertion site and flanking region. The double mutant strain was complemented by cloning the full length *sloC* or *mntH* gene into the *S. mutans* integration vector pMC340B [52] to yield plasmids pMC340B-sloC or pMC340B-mntH. The plasmids were propagated in *E. coli* DH10B and used to transform the *S. mutans* Δ*sloC*Δ*mntH* strain for integration at the *mtl* locus. All primers used in this study are listed in Table 4.

**Table 4.**
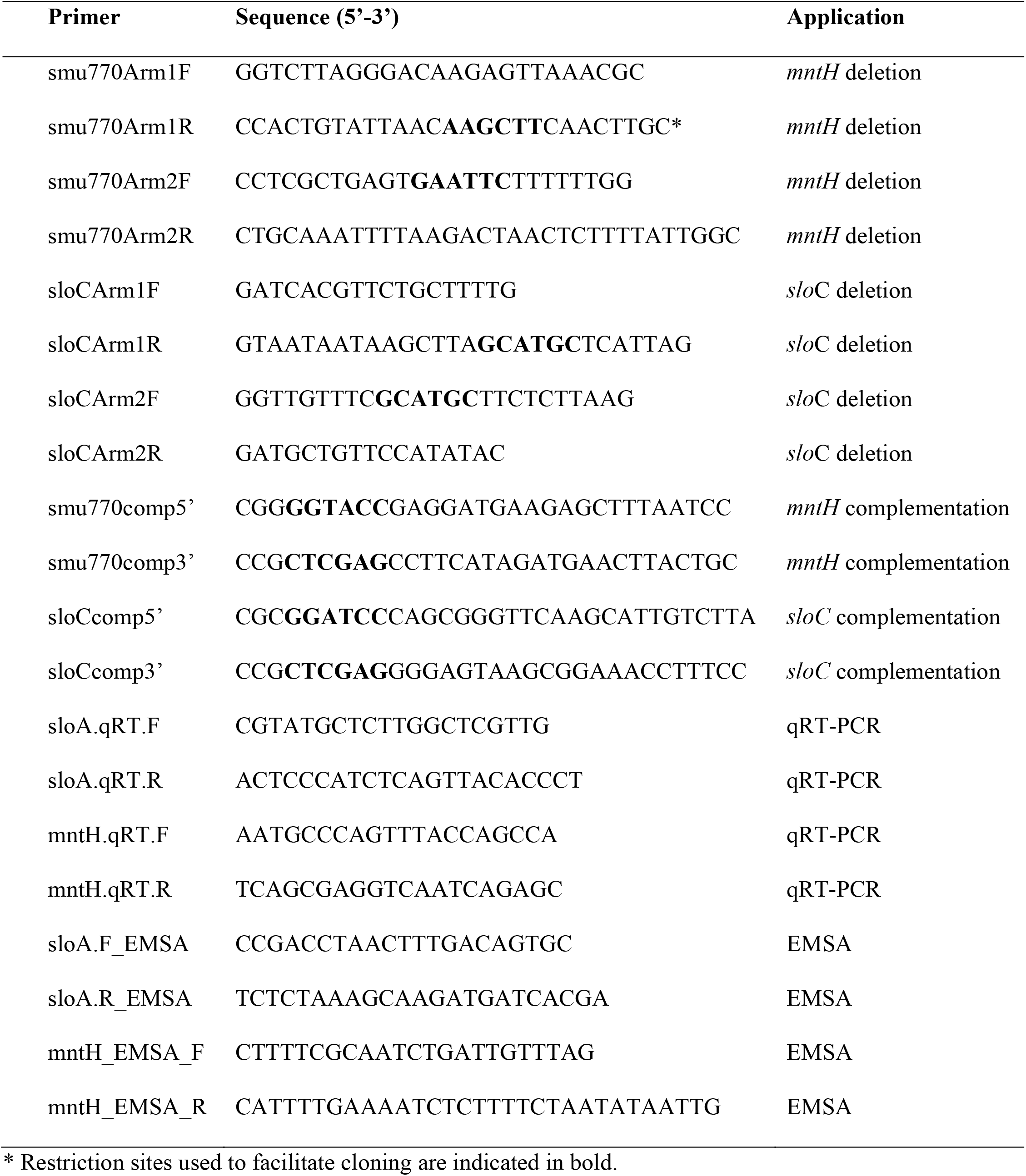
Primers used in this study.

### RNA analysis

Total RNA was isolated from homogenized *S. mutans* cell lysates by acid-phenol:chloroform extractions as previously described [53]. The RNA was precipitated with ice-cold isopropanol and 3M sodium acetate (pH 5) at 4°C before RNA pellets were resuspended in nuclease-free H_2_O and treated with DNase I (Ambion) for 30 minutes at 37°C. Then, 100 µg of RNA per sample was purified using an RNeasy Kit (Qiagen) including a second on-column DNase digestion according to the manufacturer’s instructions. Sample quality and quantity were assessed on an Agilent Bioanalyzer 2100 at the University of Florida Interdisciplinary Center for Biotechnology Research (UF-ICBR). 5 µg of RNA per sample were subjected to two rounds of mRNA enrichment using a MICROBExpress Bacterial mRNA Purification kit (ThermoFisher). cDNA libraries with unique barcodes were generated from 100 ng enriched mRNA using the NEB Next UltraII Directional RNA Library Prep kit for Illumina (New England Biolabs). The individual cDNA libraries were assessed for quality and quantity by Qubit. The cDNA libraries were then diluted to 10 nM each, and equimolar amounts were pooled together. The pooled libraries were subjected to RNA deep sequencing (RNA-Seq) at the UF-ICBR using the Illumina NextSeq500 platform. Read mapping was performed on a Galaxy server hosted by the University of Florida Research Computer using Map with Bowtie for Illumina, and the *S. mutans* UA159 genome (NC_004350.2) as a reference. The reads per open reading frame were tabulated with htseq-count. Final comparisons between the control and Mn-depleted conditions were performed with Degust (http://degust.erc.monash.edu/), with a False Discovery Rate (FDR) of 0.05, and a 2-fold change cutoff. Quantifications of *mntH* and *sloA* mRNA were obtained by quantitative real time PCR (qRT-PCR) using gene-specific primers (Table 4) on triplicate samples of the *S. mutans* UA159 and GMS584 (Δ*sloR*) strains grown to mid-logarithmic phase (OD_600_ of 0.5) according to established protocols [46]. Student’s *t*-test was applied to the analysis of the qRT-PCR results.

### ICP-OES analysis

The total metal content within bacterial cells was determined using ICP-OES performed at the University of Florida Institute of Food and Agricultural Sciences (UF-IFAS) Analytical Services Laboratories. Briefly, cultures (250 ml) were grown in plain BHI to mid-exponential phase (OD_600_ = 0.4), harvested by centrifugation at 4°C for 15 min at 4,000 RPM, and washed first in PBS supplemented with 0.2 mM EDTA to chelate extracellular divalent cations followed by a wash in PBS alone. The bacterial pellets were resuspended in 2 ml 35% HNO_3_ and digested at 90°C for 1 h in a high-density polyethylene scintillation vial. The digested bacteria were diluted 1:10 in reagent-grade H_2_O prior to ICP-OES metal analysis. The metal composition was quantified using a 5300DV ICP atomic emission spectrometer (PerkinElmer), and concentrations were determined by comparisons to a standard curve. Metal concentrations were then normalized to total protein content as determined by the bicinchoninic acid (BCA) assay (Pierce).

### Growth antagonism assay

The ability of *S. gordonii* or *S. sanguinis* to inhibit the growth of *S. mutans* via H_2_O_2_ production was assessed as described previously [54, 55]. Briefly, 8 µl of an overnight culture of *S. gordonii* DL-1 or *S. sanguinis* SK150 was spotted in the center of a BHI+Mn agar plate and incubated at 37°C, 5% CO_2_. After 24 hr incubation, 8 µl of *S. mutans* overnight cultures grown in BHI+Mn were spotted near the peroxigenic strain, and similarly allowed to incubate overnight before monitoring for proximal growth defects. To confirm that growth inhibition was due to H_2_O_2_ production, a control condition was included in which 8 µl of 1mg ml^-1^ catalase solution was spotted on top of the peroxigenic strain spot prior to spotting the *S. mutans* culture.

### Growth and survival in human saliva

To test the ability of the *S. mutans* strains to proliferate and survive in saliva, pooled human saliva was filter-sterilized using a 0.2 micron membrane and heat-inactivated at 65°C for 30 min. Cultures of *S. mutans* grown in BHI to an OD_600_ of 0.25 as described above were then diluted 1:20 into filtered saliva supplemented with either 10 mM glucose, or with 10 mM glucose and 10 µM MnSO_4_ prior to incubation at 37°C in a 5% CO_2_ atmosphere. Immediately upon dilution in saliva and at selected time intervals, 10-fold serial dilutions were prepared in sterile PBS and plated onto BHI+Mn agar for viable plate counting. Saliva samples were collected after obtaining written consent as per the study approval from the University of Florida Internal Review Board (Protocol #: 201600877).

### Biofilm assay

The ability of *S. mutans* strains to form biofilms on saliva-coated wells of polystyrene microtiter plates was assessed by growing cells in BHI supplemented with 1% sucrose with or without supplementation with 10 μM of Mn. The wells of the plates were first coated for 30 minutes with 100 μl of sterile, clarified, pooled human saliva. Next, strains grown in BHI+Mn to an OD_600_ of 0.5 were diluted 1:100 in BHI containing 1% sucrose, and added to the wells of the microtiter plate. Plates were incubated at 37°C in a 5% CO_2_ atmosphere for 4 and 24 h. After incubation, plates were washed twice with water to remove planktonic and loosely-bound bacteria, and adherent cells were stained with 0.1% crystal violet for 15 min. The bound dye was eluted with 33% acetic acid solution, and biofilm formation was then quantified by measuring the optical density of the solution at 575 nm.

### Electrophoretic mobility shift assay

EMSAs were performed according to established protocols [46]. Briefly, primers were designed to amplify the promoter regions of the *S. mutans sloABC* and *mntH* genes (Table 4). The resulting amplicons were end-labeled with [γ-^32^P] dATP (Perkin-Elmer) in the presence of T4 polynucleotide kinase (New England BioLabs), after which they were centrifuged through a TE Select-D G-25 spin column (Roche Applied Science) to remove unincorporated [^32^P] dATP. Binding reaction mixtures were prepared as 16 µl reactions containing 1 µl (∼13.25ng) of end-labeled amplicon, purified native SloR protein at concentrations ranging from 0 to 400nM, and 3.2 µl of 5x binding buffer (42 mM NaH_2_PO_4_, 58 mM Na_2_HPO_4_, 250 mM NaCl, 25 mM MgCl_2_, 50 mg ml^-1^ bovine serum albumin, 1 mg sonicated salmon sperm DNA, 50% glycerol, and 37.5 M MnCl_2_). Samples were loaded onto 12% nondenaturing polyacrylamide gels and resolved at 300 V for 1.5 h. Gels were exposed to Kodak BioMax film for 24 h at 80°C in the presence of an intensifying screen prior to autoradiography.

## Data availability

Gene expression data have been deposited in the NCBI Gene Expression Omnibus (GEO) database (www.ncbi.nlm.nih.gov/geo) under GEO Series Accession number GSE139093.

## Acknowledgements

This study was supported by NIH-NIDCR R01 DE019783 and NIH-NIAID R21 AI137446 to JAL and NIH-NIDCR R01 DE014711 to GAS. Purified calprotectin was generously provided by Eric Skaar and Walter Chazin at Vanderbilt University.

